# Leveraging HIV-Specific CAR T Cells and Rapamycin Treatment in “Kick-and-Kill” HIV Cure Approaches

**DOI:** 10.64898/2026.01.15.699756

**Authors:** Nishad S. Maggirwar, José A. Morán, Wenli Mu, Thomas D. Zaikos, Tessa Chou, Shireen R. Turner, Brian H. Yu, Alok Ranjan, Rami Hourani, Paul A. Wender, Anjie Zhen, Matthew D. Marsden

**Affiliations:** Department of Microbiology and Molecular Genetics, School of Medicine, University of California Irvine; Department of Medicine, Division of Hematology and Oncology, University of California Los Angeles; AIDS institute, University of California Los Angeles; Department of Chemistry, Stanford University; Department of Chemical and Systems Biology, Stanford University; Department of Medicine, Division of Infectious Diseases, School of Medicine, University of California Irvine

## Abstract

HIV is not cured with currently available combination antiretroviral therapy (ART) alone in large part because the virus establishes virologic latency in long lived CD4^+^ cells. To eliminate this latent reservoir, as required for HIV eradication, latency reversing agents (LRAs) are being developed to force HIV out of latency and induce infected cells to express viral proteins leading to their clearance, in a so-called “Kick-and-Kill” approach. This strategy relies on the immune system to clear the productively-infected cells and is thus limited by HIV immune evasion and the immunological exhaustion that occurs during HIV infection. To counter these limitations and augment an LRA-mediated HIV cure approach, we report herein the utility of HIV-specific truncated CD4-based D1D2 CAR T cells combined with LRA treatment and the mTORC1 inhibitor rapamycin to reduce immune exhaustion and specifically target and kill LRA-stimulated HIV infected cells. We demonstrate that rapamycin does not prevent HIV latency reversal via multiple classes of LRAs in several *in vitro* models, suggesting that it is compatible with cure approaches utilizing these LRAs. Additionally, rapamycin does not inhibit the early T cell activation (CD69 upregulation) in primary T cells that occurs during protein kinase C (PKC) modulator-mediated HIV latency reversal. Furthermore, *in vitro* chronically exhausted CAR T cells were found to have a higher frequency of terminally exhausted PD-1^+^Tim-3^+^ and CD69^+^PD-1^+^ cells when compared to CAR T cells that were cultured under the same conditions in the presence of rapamycin, validating the use of the mTORC1 inhibitor rapamycin to prevent immune exhaustion of CAR T cells. Finally, we found that latently-infected cells that were stimulated to express HIV proteins using a designed, synthetic PKC modulator LRA (SUW133) were efficiently recognized and killed by CAR T cells. Overall, these data demonstrate the compatibility of immune rejuvenation using rapamycin with HIV reservoir depletion using LRAs and CAR T cells. This combination therapy strategy represents a promising approach to more effectively target the latent reservoir in HIV cure approaches.

## INTRODUCTION

A scalable, functional cure for HIV currently does not exist due to several factors including the chronic nature of HIV infection, error-prone viral replication machinery, and the establishment of viral latency within long lived CD4^+^ cells(1–4). Consequently, people with HIV (PWH) who are treated with antiretroviral therapy (ART) to prevent disease progression still harbor persistent latent reservoirs of virus that would resupply virus if ART is stopped.

Several approaches have been undertaken to address these barriers to developing a safe and effective HIV cure. One experimental HIV eradication strategy known as the “Kick-and-Kill” approach involves forcing the virus to express viral proteins in infected CD4^+^ cells using latency reversing agents (LRAs) and allowing immune clearance mechanisms or viral cytopathic effects to kill the infected cells(5–8). Multiple classes of LRAs with distinct mechanisms of reactivating HIV infected cells have been identified, including but not limited to, protein kinase C (PKC) modulators(6,9,10), second mitochondrial-derived activator of caspases (SMAC) mimetics(11–13), histone deacetylase inhibitors (HDACis)(14–17), and bromodomain extra-terminal motif (BET) bromodomain inhibitors(18–20). PKC modulators compete with the endogenous diacylglycerol ligand for activation of PKC which in turn targets the NF-kB pathway and triggers downstream signaling pathways that induce NF-kB-mediated transcription of latent HIV(6,21). HDACis remove epigenetic blocks that otherwise silence viral expression by preventing chromatin deacetylation, thereby promoting chromatin accessibility by transcription factors(22,23). Finally, BET-bromodomain inhibitors bind to BET bromodomain protein BRD4, which competes with HIV Tat for access to the host transcription factor p-TEFb, preventing BRD4 – p-TEFb interaction, and promoting HIV Tat-mediated viral transcription(19).

While this kick-and-kill approach has shown promise in reversing HIV from latency and reducing rebound-competent reservoirs in various preclinical studies(6,24,25), further advances are needed to completely deplete the latent reservoir and develop a cure(26). This includes applying methods to ensure that effective HIV-specific cellular immune responses are present to kill latently-infected cells when they are induced to express viral proteins. This killing of infected cells can be obstructed by factors including immune evasion mechanisms by HIV, immune exhaustion, the immunological damage that is a hallmark of HIV infection, and the waning HIV-specific immune response that often occurs during long-term ART when viral antigens are scarce(26).

One potentially powerful approach to overcome these issues is the use of HIV-specific chimeric antigen receptor (CAR)-expressing T cells, which have been genetically modified to express surface receptors that recognize HIV infected cells. One such CAR cell approach utilizes D1D2CAR 4-1BB(27), encoding a truncated version of CD4 that can bind to cells expressing HIV *env*, which signals the CAR cell to kill the infected target cell. This CD4-gp120 interaction is difficult for HIV to evolve around, and the truncated receptor consisting of only D1 and D2 domains of CD4 (with D3 and D4 domains absent) make this a poor entry receptor for HIV. Moreover, this interaction is MHC-unrestricted(27,28) which circumvents the necessity for a TCR:MHC interaction, and therefore presentation of mutated HIV MHC I peptides will not interfere with CAR T cell recognition and killing of infected cells. Finally, the vector expressing this CAR encodes shRNAs against CCR5 and the HIV 5’LTR, which act as safety features and further limit the ability of these cells to support productive infection by HIV(28–30).

While CAR T cells are potentially powerful tools for use in HIV cure approaches, in the context of chronic infections such as HIV, immune cells including CAR T cells can become immunologically exhausted and impaired in their ability to clear infected cells(4,28,31). Terminally exhausted CAR T cells (and other effector cells such as CD8^+^ T cells), are defined by those that are PD-1^+^Tim-3^+^ and upregulating other markers such as CD69 and CD39(32–34). These exhausted cells show decreased proliferative capacity, as well as decreased production of key inflammatory cytokines such as TNF, IL-2, and IFNγ, ultimately resulting in a failure to clear infected cells(35). In this study, we have investigated the utility of rapamycin to directly decrease the frequency of terminally exhausted PD-1^+^Tim-3^+^ and PD-1^+^CD69^+^ cells during chronic *in vitro* stimulation, to mimic conditions that would be observed during a chronic HIV infection. Previously, rapamycin has been shown to reduce immune cell exhaustion via the inhibition of mammalian target of rapamycin complex 1 (mTORC1), to improve mitochondrial function, and to improve cytokine production, resulting in enhanced effector cell function(4,28,36).

HIV-specific CAR T cells have been used in combination with low and intermittent treatment with rapamycin, which led to delay in viral rebound and a decrease in viral load upon ART cessation in humanized mouse models infected with HIV(4,27,28). Here, we explore whether a similar CAR T cell and rapamycin approach is compatible with the use of HIV latency reversal agents (LRAs) to purge latently-infected cells. We show that rapamycin does not interfere with HIV latency reversal using several different classes of LRAs, that rapamycin rejuvenates *in vitro* exhausted primary CAR T cells, and that a promising designed, synthetic, protein kinase C modulator can induce expression of latent HIV and make the target cells susceptible to CAR T cell-mediated killing when administered with rapamycin. Hence, a combination chemotherapy approach using rapamycin-rejuvenated CAR T cell and LRAs represents a potentially useful approach to HIV reservoir depletion.

## RESULTS

### Rapamycin inhibits primary human peripheral blood mononuclear cell (PBMC) proliferation during αCD3/αCD28 co-stimulation

The immunosuppressive effects of rapamycin on inflammation, immune cell activation and proliferation have been well documented in the literature(4,37–39). To verify the findings previously described in other relevant experimental systems(40,41), we isolated primary human PBMC from healthy donors and co-stimulated them by ligating CD3 and CD28 in the presence of IL-2, and treated with 5 nM or 5000 nM rapamycin. This resulted in decreased proliferative capacity compared to PBMCs that were stimulated in the absence of rapamycin **(Fig. 1)**. This is consistent with rapamycin’s ability to inhibit the transition of G1 to S in proliferating immune cells(38).

**Fig. 1.**
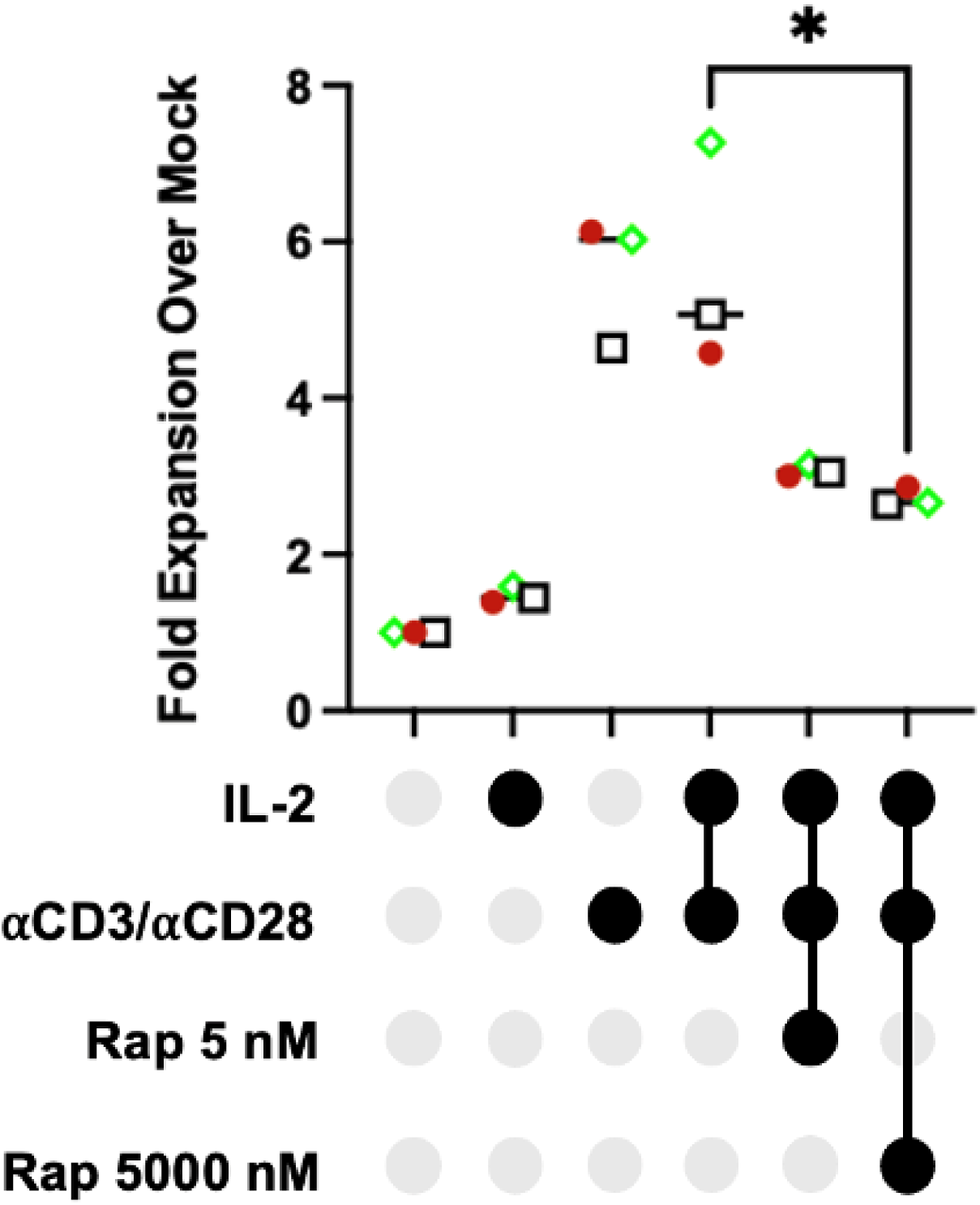
Rapamycin inhibits PBMC proliferation during co-stimulation. PBMCs were partitioned into wells and treated with a combination of 30 U/mL IL-2, ⍺CD3/⍺CD28 magnetic beads, and rapamycin (5 nM or 5000 nM). Cells were incubated for 4 days and quantified. Significance assigned by unpaired t-test. *, P = 0.02. N = 3 unique HIV seronegative human PBMC donors (indicated by a unique colored symbol).

### Rapamycin is compatible with HIV latency reversal in J-Lat 10.6 and A2 clones using 3 different classes of LRAs

Having verified that rapamycin is functioning as expected in a primary human PBMC proliferation assay, we then evaluated its effects on HIV latency reversal in relevant cell line models to assess whether it is compatible for use with LRAs. To quantify this, the latently-infected Jurkat-Latency (J-Lat) clones 10.6 and A2 were treated with several relevant doses of LRAs and rapamycin. J-Lat cell lines harbor a well characterized, replication incompetent, latent HIV provirus that encodes green fluorescent protein (GFP). This allows measurement of HIV latency reversal via GFP quantification, without the added complications of HIV replication and spread within an *in vitro* culture. J-Lat 10.6 clones contain a near full length HIV provirus with defective *env* and *nef* genes, while J-Lat A2 clones contain a truncated HIV provirus solely consisting of HIV *tat* (HIV transcriptional activator required for HIV expression), an internal ribosomal entry site (IRES), and a GFP gene(42,43). Because J-Lat 10.6 and A2 clones primarily differ in integration site and HIV viral gene content, we were able to observe how these variables contribute to HIV latency reversal *in vitro* in combination with LRAs and rapamycin. Compound dosing was determined based on prior studies using these LRAs(6,10,14,16,23,44,45). We observed that rapamycin, when used in combination with three different classes of LRAs including PKC modulators (bryostatin-1 and SUW133, **Fig. 2A and Fig. 2B**), the BET bromodomain inhibitor JQ1(+) **(Fig. 2C)**, and several HDAC inhibitors (panobinostat, entinostat, or vorinostat) (**Fig. 2D**), exhibits in some cases a slight antagonistic effect on HIV latency reversal, but does not prevent it. Additionally, we observed that SUW133 induced the greatest levels of latency reversal with or without rapamycin in both J-Lat cell line clones. This result was expected, as SUW133 is a synthetic analog of one of the strongest LRAs *in vitro*, bryostatin-1, and has been shown to induce stronger latency reversal *in vitro* and *in vivo*, compared to bryostatin-1(6,24,25). When SUW133 was combined with JQ1(+) and rapamycin, almost all J-Lat cells in the culture were induced to express HIV. JQ1(+) has been shown to function synergistically with other drugs in the context of cancer and HIV(18,46–49). Based on the Bliss Independence Model(50), JQ1(+) synergized with SUW133 and rapamycin to induce greater latency reversal in both J-Lat 10.6 and A2 clones. We also observed that PKC modulators (SUW133 and bryostatin-1) outcompeted HDAC inhibitors in the amount of latency reversal they induced in J-Lat cell lines. Rapamycin, when used at 5 nM had a significant effect on latency reversal induced by panobinostat (50 nM) and entinostat (1000 nM), but ultimately did not prevent it **(Fig. 2D)**.

**Fig. 2.**
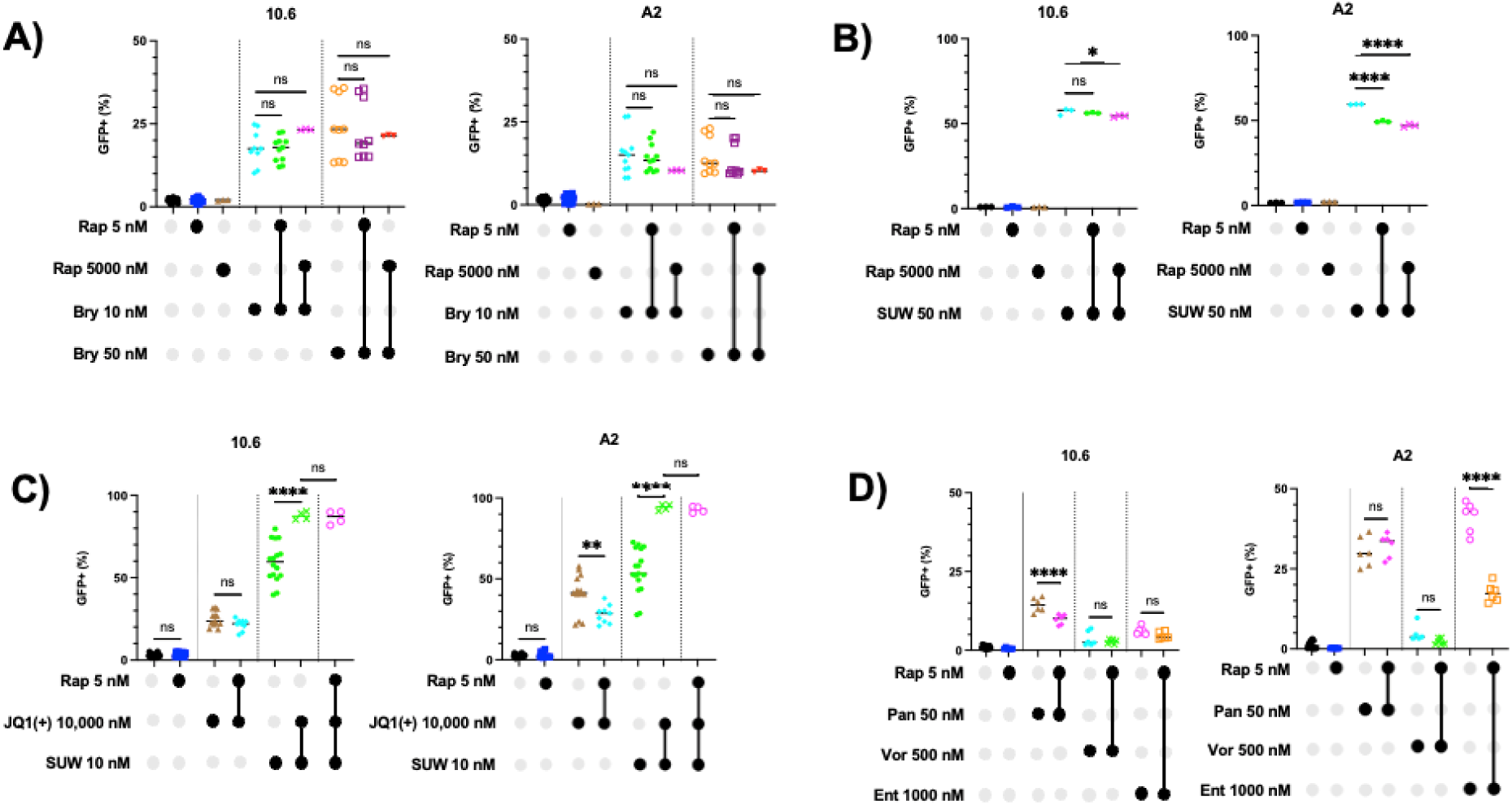
Rapamycin does not prevent HIV latency reversal induced by various classes of LRAs in J-Lat clones 10.6 and A2. A-D) J-Lat clones 10.6 and A2 were treated with various classes of latency reversing agents and rapamycin. Extent of latency reversal was measured via GFP+% expression using flow cytometry. Rap=Rapamycin, SUW=SUW133, JQ1(+)=JQ1(+), Bry=Bryostatin-1, Pan=Panobinostat, Vor=Vorinostat, Ent=Entinostat. Shaded circles indicate treatment conditions. Significance assigned by one-way ANOVA with Tukey’s multiple comparisons test. *, P = 0.02; **, P = 0.005; ****, P < 0.0001; ns, P > 0.05. N = 3-31 technical replicates per treatment condition.

### Rapamycin does not inhibit immune activation markers upon LRA treatment or co-stimulation in T lymphocytes

One biomarker on T cells that has been correlated with HIV latency reversal *in vivo* in the context of PKC modulators and co-stimulatory signaling is the early T lymphocyte activation marker CD69(6,10). To evaluate the effects of rapamycin on early and late T cell activation and determine whether the reduction in PBMC proliferation observed (**Fig. 1)** was caused by a decrease in CD25, we co-stimulated bulk human PBMCs in the presence of IL-2 with or without rapamycin, and quantified CD69 (early T cell activation) and CD25 (late T cell activation) expression on T lymphocytes via flow cytometry. CD25 (also known as IL-2Rα) is the alpha-chain of the IL-2 receptor heterotrimer, and upon binding to IL-2 induces signaling that promotes T cell proliferation(51). We observed that co-stimulated PBMCs that were also treated with rapamycin did not show decreased CD69 expression on CD4^+^ **(Fig. 3A)** and CD8^+^ T lymphocytes **(Fig. 3C)**. However, CD4^+^ T lymphocytes **(Fig. 3B)** and CD8^+^ T lymphocytes **(Fig. 3D)**, did show a modest decrease in CD25 expression in the 5000 nM rapamycin treatment condition.

**Fig. 3.**
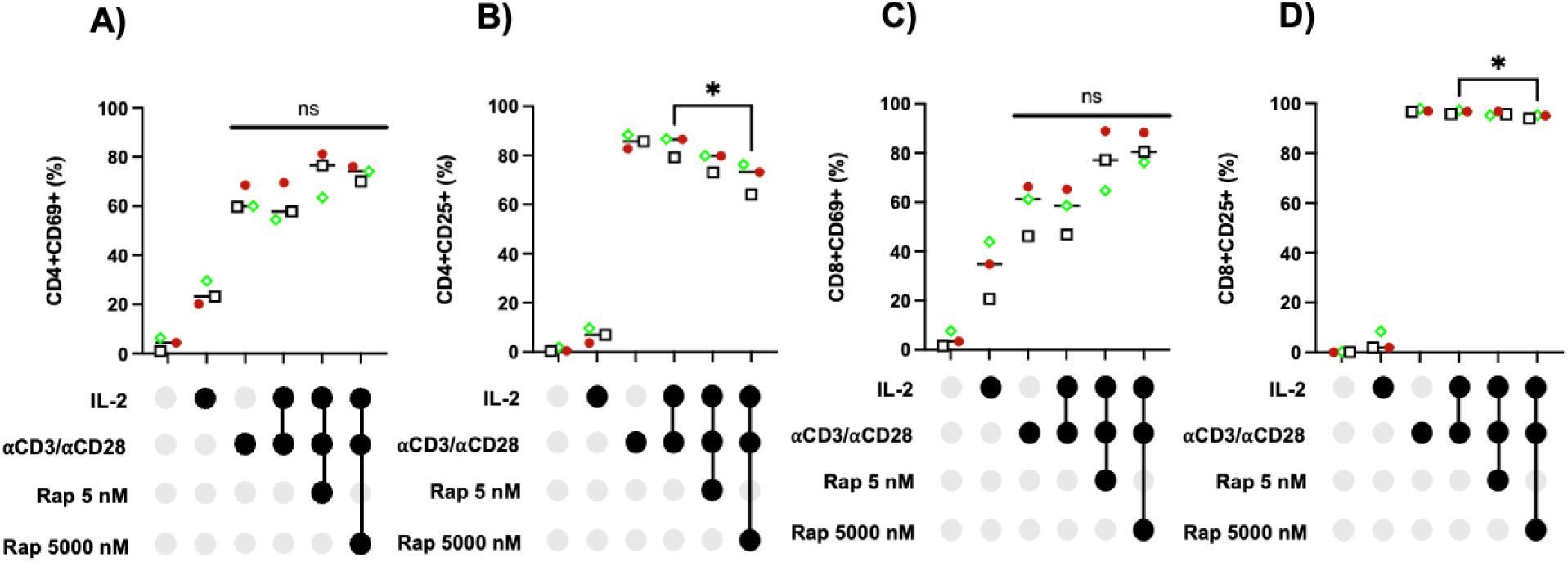
Rapamycin does not prevent early and late T cell activation markers during co-stimulation. PBMCs were treated with 30 U/mL IL-2, ⍺CD3/⍺CD28 magnetic beads, 5 nM or 5000 nM rapamycin. After 4 days of incubation, PBMCs were assessed for A and C) CD69 and B-D) CD25 expression via flow cytometry. Significance assigned by one-way ANOVA with Tukey’s multiple comparisons test. *, P = 0.04; ns, P > 0.05. N = 3 unique HIV seronegative human PBMC donors (indicated by a unique colored symbol).

We also evaluated the effect of rapamycin on CD69 expression in bryostatin-1-treated bulk human PBMCs, which were then analyzed via flow cytometry to measure CD69 expression on CD4^+^ T lymphocytes **(Fig. 4A)** and CD8^+^ T lymphocytes **(Fig. 4B)**.

**Fig. 4.**
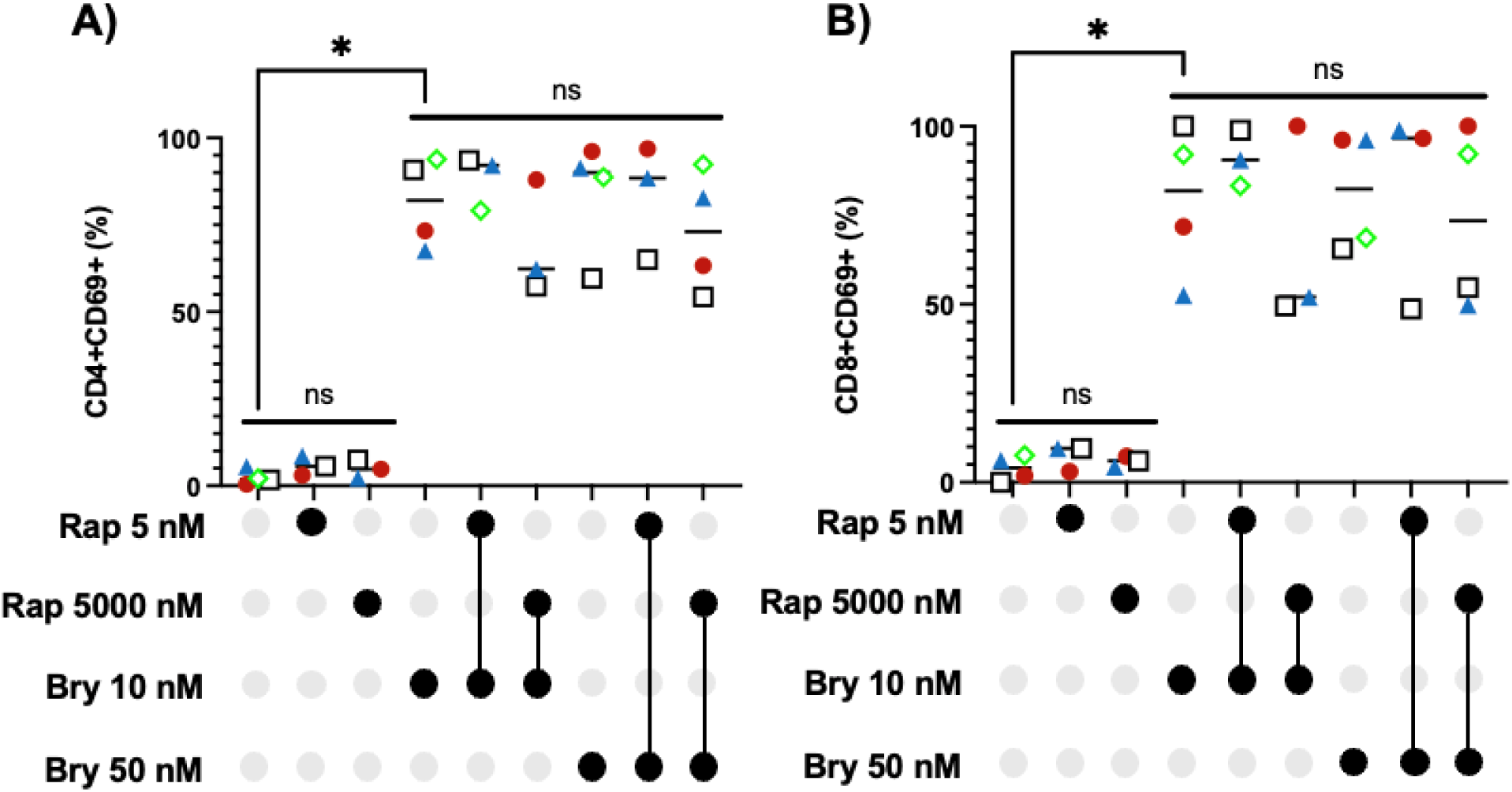
Rapamycin does not prevent early T cell activation during bryostatin-1 treatment. PBMCs were isolated from human blood and stained for markers of T lymphocyte activation. A) CD4^+^CD69^+^ and B) CD8^+^CD69^+^ cell populations were identified via flow cytometry. Significance assigned by one-way ANOVA with Tukey’s multiple comparisons test. *, P = 0.01; ns, P > 0.05. N = 3-4 unique HIV seronegative human PBMC donors (indicated by a unique colored symbol).

Similar to the co-stimulation condition (**Fig. 3A-D**), rapamycin, when used at the indicated concentrations, did not decrease CD69 expression in CD4^+^ and CD8^+^ T lymphocytes. This data indicates that rapamycin does not interfere with the early T cell activation pathways that are correlated with successful HIV latency reversal in the context of PKC modulators such as bryostatin-1 and SUW133(6), which provides encouraging support for a “Kick-and-Kill” HIV cure approach that incorporates rapamycin.

### HIV-specific CAR T cells recognize and specifically kill latently-infected cells treated with the designed, synthetic PKC modulator SUW133

To determine whether CAR T cells can recognize latently infected target cells treated with the designed synthetic LRA SUW133, CAR T cells were generated *in vitro* by lentiviral transduction of primary human CD8^+^ T cells, and co-cultured with promonocytic U1 cells that had been pre-treated with SUW133. U1 cells harbor two HIV proviruses with mutated *tat* genes, one containing an ATG initiation codon, and another containing an H_13_èL mutation(52–54). Hence, they require a strong stimulus to induce viral protein expression in the absence of Tat, and once activated by LRAs, they express HIV Env on their surface, which serves as the target for D1D2 CAR T cells. We also confirmed that our CAR construct expresses a truncated CD4, as evidenced by the selective binding with a D1-specific antibody and lack of binding with an antibody specific to the D3 subunit of CD4 **(Fig. S1)**. We observed that CAR T cells incubated with SUW133-stimulated U1 cells showed increased expression of CD107a on the CAR T cells, compared to the mock condition **(Fig 5)**. In this co-incubation model, anti-CD107a antibody was added directly to the culture in the presence of the monensin-containing protein transport inhibitor GolgiStop, to maximize staining of CD107a which is upregulated in response to effector cell degranulation, and re-internalized(55,56). Increased expression of CD107a on CAR T cells in the presence of SUW133-activated U1 cells indicated effector cell specific recognition of target cells. Furthermore, this observation indicates CAR T cell recognition of U1 cells that have been reversed from latency using SUW133 **(Fig. 5A and Fig. 5B)**. To measure specific killing of SUW133-activated U1 cells by CAR T cells, CAR T cells and SUW133-treated U1 cells were co-incubated as previously described. Expression of the HIV polyprotein Gag (comprised of the viral structural proteins p17 matrix (MA), p24 capsid (CA), p7 nucleocapsid (NC), and p6(57)) is indicative of a productively infected cell, which can then be selectively identified by CAR T cells, as productively infected cells will likely express HIV *env*. We observed greater specific killing of U1 cells treated with SUW133 compared to non-SUW133 treated U1 cells, indicating that CAR T cells can effectively kill U1 cells that have been induced to express HIV using the novel LRA, SUW133 **(Fig. 6A and Fig. 6B)**.

**Fig. 5.**
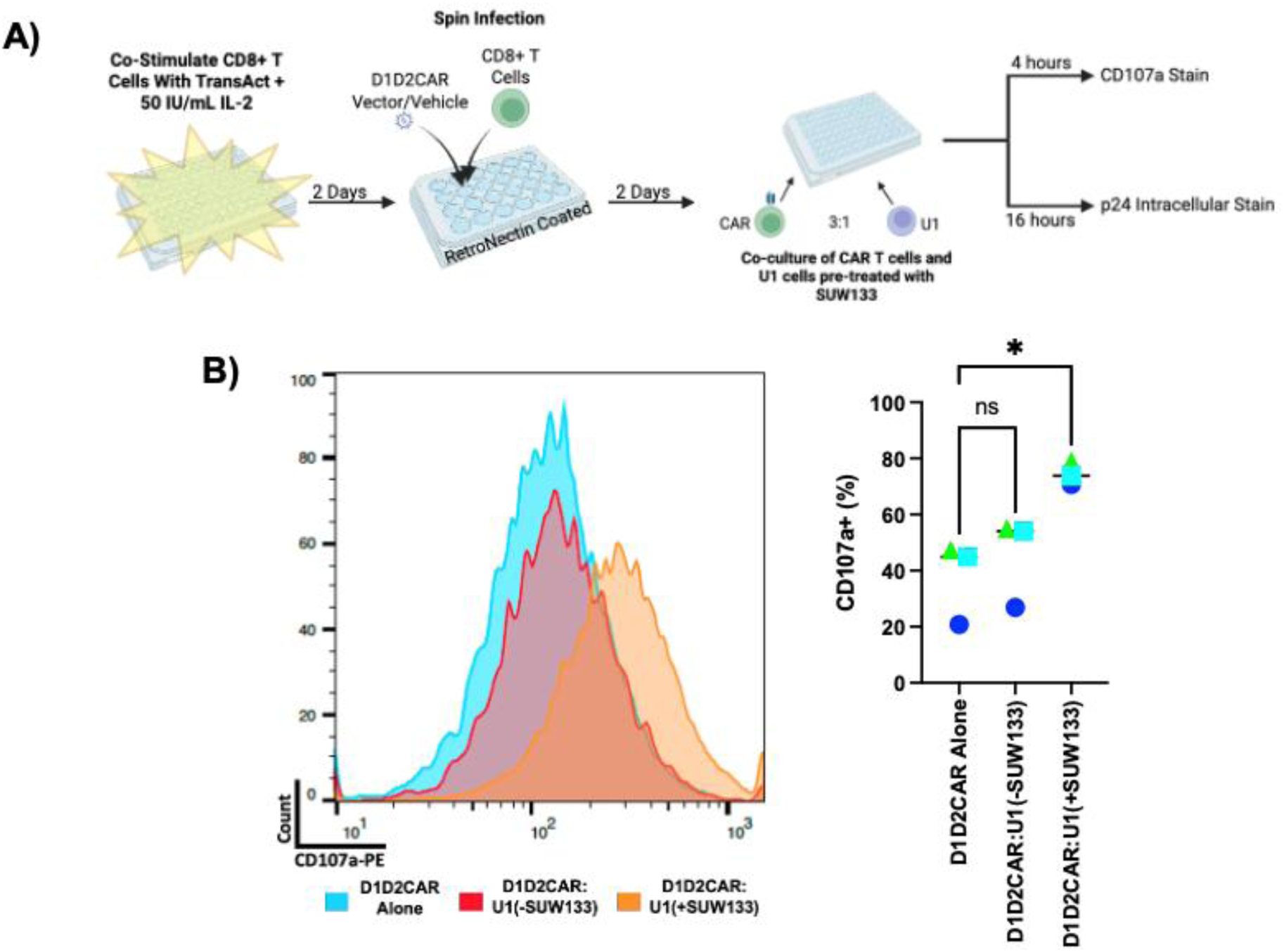
Greater CD107a expression is observed when D1D2CAR T cells are co-incubated with U1 cells treated with SUW133. A) D1D2CAR T cells were generated *in vitro* via D1D2CAR 4-1BB lentiviral transduction of primary human CD8^+^ T cells and cultured in growth media containing 50 IU/mL IL-2 for one week. U1 cells were mock treated (-SUW133) or treated with 50 nM SUW133 (+SUW133) for two days. D1D2CAR 4-1BB CAR T cells were mixed at a 3:1 effector to target ratio with mock-treated U1 cells (D1D2CAR:U1(-SUW133)) or 50 nM SUW133-treated U1 cells (D1D2CAR:U1(+SUW133)). D1D2CAR 4-1BB CAR T cells were also assessed alone to determine baseline CD107a levels. B) CD107a expression was quantified. Significance assigned by paired t-test. *, P = 0.03; ns, P > 0.05. N = 3 unique HIV seronegative human PBMC donors (indicated by a unique colored symbol).

**Fig. 6.**
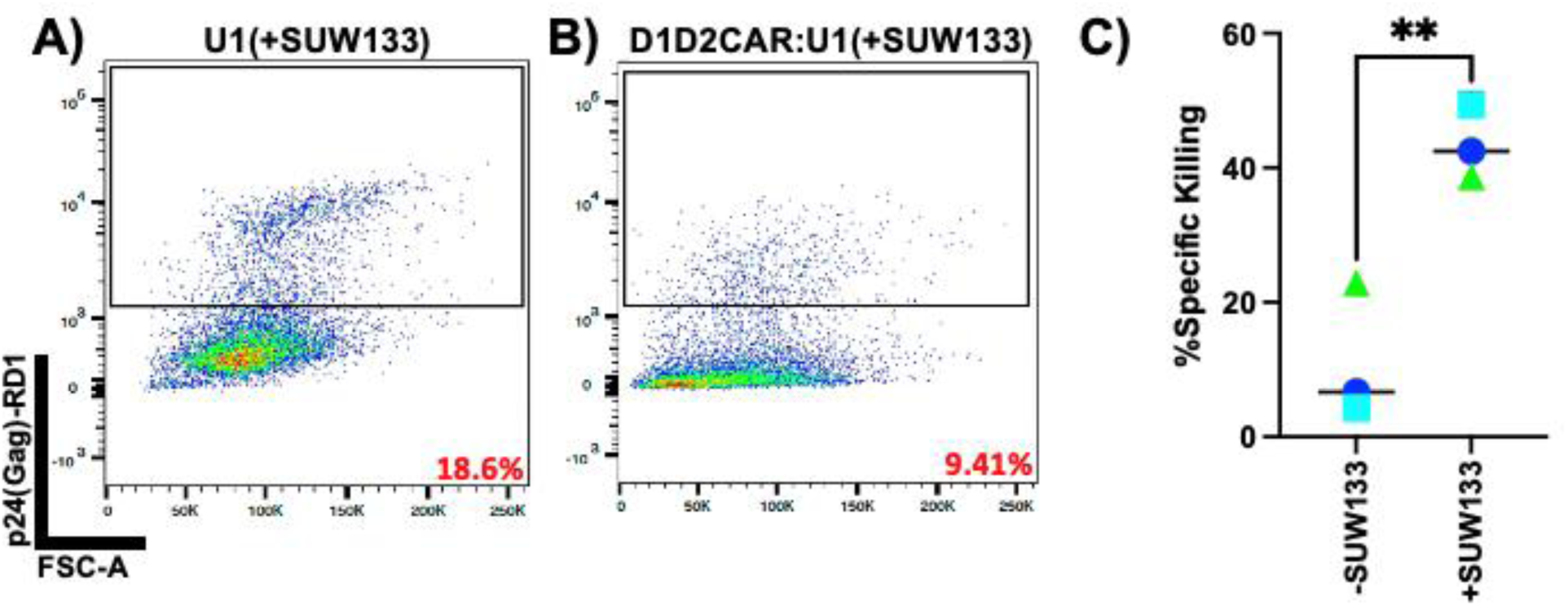
D1D2CAR T Cells Selectively Kill U1 Cells Treated with SUW133. D1D2CAR T cells were generated *in vitro* and co-incubated with mock (-SUW133) or A) SUW133 treated (+SUW133) U1 cells at a 3:1 effector to target ratio. B) D1D2CAR T cells and U1 cells were incubated for 16 hours before assessing p24 expression via flow cytometry. D) D1D2CAR T cell-specific killing of SUW133-treated U1 cells in multiple human PBMC donors. Significance assigned by unpaired t-test. **, P = 0.0084. N = 3 unique HIV seronegative human PBMC donors (indicated by a unique colored symbol).

### Rapamycin treated HIV-specific CAR T cells show a decreased frequency of terminal immune exhaustion markers

CAR T cell exhaustion in the context of chronic infections, such as HIV, is a critical barrier to developing an HIV cure(28,31). We sought to determine whether rapamycin can decrease the frequency of terminally exhausted CAR T cells *in vitro*. After generating CAR T cells *in vitro,* CAR T cells were co-stimulated for two days, followed by 5 days of αCD3 treatment. This consistent αCD3 treatment enforces clustering of the CD3 co-receptor, leading to antigen-independent chronic stimulation(58). We observed that there was no statistically significance difference in the percentage of PD-1^+^Tim-3^+^ cells between unstimulated CAR T cells and cells that were chronically stimulated in the presence of rapamycin **(Fig. 7A-B)**. Consistent with **Fig. 4** and **Fig. 5**, rapamycin did not induce a significant decrease in immune activation markers CD69 and CD25 upon stimulation. Additionally, rapamycin-treated chronically-stimulated CAR T cells showed a decreased percentage of PD-1^+^CD69^+^ cells **(Fig. S2)**, further suggesting that rapamycin can decrease CAR T cell exhaustion markers *in vitro*. Together these data show that CAR T cells have decreased expression level of PD-1 and Tim-3 in the presence of rapamycin during chronic stimulation conditions yet continue to stay immunologically active.

**Fig. 7.**
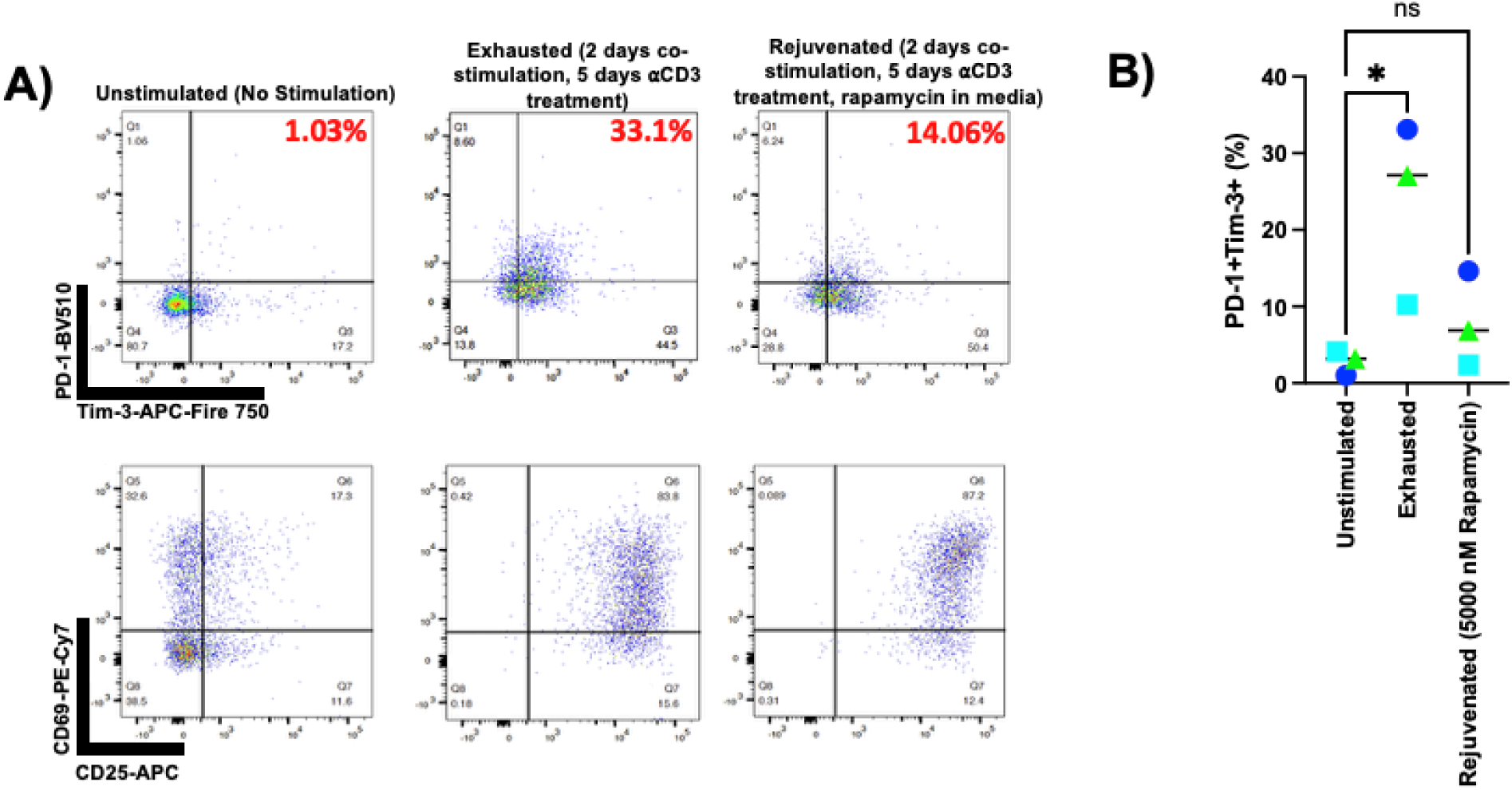
PD-1^+^Tim-3^+^ cells are less abundant in chronically stimulated, rapamycin treated, D1D2CAR T cells. A) D1D2CAR T cells were generated *in vitro* and co-stimulated in ɑCD3 coated plates with ɑCD28 and 50 IU/mL IL-2 in growth media. After two days of co-stimulation, cells were placed into new ɑCD3 coated plates, without ɑCD28 but with 50 IU/mL IL-2. Every two days, cells were moved to a freshly ɑCD3-coated plate with fresh growth media. After 7 days, cells were collected and assessed for expression of B) immune exhaustion markers and immune activation markers via flow cytometry. C) %PD-1^+^Tim-3^+^ double positive cells were quantified. Significance assigned by one-way ANOVA with Tukey’s multiple comparisons test. *, P = 0.04; ns, P > 0.05. N = 3 unique HIV seronegative human PBMC donors (indicated by a unique colored symbol).

## DISCUSSION

HIV can evade immune detection and clearance through an array of different mechanisms. These include selective MHC I downregulation and error-prone reverse transcription, leading to the presentation of mutated viral antigens that evade CTL responses(29,59). Coupled with the propensity of the virus to form long-lived latent reservoirs in CD4^+^ T cells that persist for life even during ART, these mechanisms represent formidable challenges to curing HIV infection. Hence, to develop a scalable, effective HIV cure approach, a multi-pronged strategy is likely necessary. Here, we have explored the combination of latency reversing agents, HIV-specific CAR T cells, and mTORC1 inhibitor rapamycin as a strategy to significantly enhance HIV-specific immunity and eliminate latently-infected cells.

The ”Kick-and-Kill” approach has shown promise as an HIV cure strategy that can eradicate latent HIV reservoir cells, and delay viral rebound after ART cessation in small animal preclinical models(6,24,25). However, this approach is limited by several potential barriers. In addition to the capacity of HIV to evade the immune response, the chronic nature of HIV infection and associated persistent immune activation leads to exhaustion of immune effector cells(3,4,28). As a result of this immune exhaustion, LRA-activated cells that are expressing viral proteins may be incompletely cleared, leading to persistence of the infection(60,61). Rapamycin has shown promise in rejuvenating exhausted immune cells in various pre-clinical models and human patient studies in the context of chronic infections and cancer(3,4,28,36,39). Additionally, as an mTORC1 inhibitor and autophagy inducer, rapamycin can exert immunomodulatory effects on immune cells, such as G1/S cell cycle arrest and a decrease in the release of several pro-inflammatory cytokines including TNFα, IL-2 and IFNγ(38,39,62,63).

Because HIV expression is connected to the activation state of the host cell, and cellular activation is also important for effector cell function, we first sought to observe whether rapamycin would interfere with either HIV latency reversal or immune cell activation.

Rapamycin decreased proliferation of co-stimulated PBMCs as expected from prior studies(4,37–39) **(Fig. 1)** but it did not prevent HIV latency reversal in J-Lat 10.6 and A2 clones that had been treated with various classes of LRAs (BET bromodomain inhibitors, HDAC inhibitors, and PKC modulators) and rapamycin **(Fig. 2A-D)**. Hence, while rapamycin has been shown to affect cell proliferation, it did not prevent HIV latency reversal via multiple individual or combination LRA approaches. It is also important to note that the doses of rapamycin that are required in the context of HIV “Kick-and-Kill” approaches, including the assays performed here and in previous publications (PMID: 39932788, 36509289) are low-dose and short-duration/intermittent. Prolonged and high-dose treatment with rapamycin as used for immunosuppression in allograft transplantation(64–66) would be counterproductive and potentially lead to ineffective killing of LRA-activated cells. The lack of HIV latency reversal inhibition in J-Lat 10.6 and A2 clones treated with LRAs and rapamycin was consistent with the lack of CD69 and CD25 downregulation in co-stimulated, and rapamycin treated PBMCs **(Fig. 3A-D)**, as well as PBMCs treated with bryostatin-1 and rapamycin **(Fig. 4A and Fig. 4B).** These data demonstrate that rapamycin does not prevent HIV latency reversal *in vitro*, and does not compromise early T cell activation pathways in a way that would significantly interfere with immune cell function, as indicated by a continued upregulation of CD69 in PBMCs in the presence of rapamycin, thus providing compelling evidence that rapamycin will not hinder our HIV cure approach.

A highly promising approach to enhance immunity against HIV is the use of HIV-specific CAR T cells. These CAR T cells can continue to recognize and kill infected cells that harbor virus with immune escape mutations that would evade recognition from the prevailing CTL responses(61,67). This is because our CAR T cells exploit the MHC unrestricted interaction between the D1 subunit of a truncated CD4 **(Fig. S1)** and HIV gp120, an interaction that is difficult for HIV to mutate around while retaining infectivity and preventing infection of the CAR T cell itself, as the CD4 truncation prevents HIV engagement of its entry co-receptors CXCR4 and CCR5(27,28). As added safety features to prevent infection of the CAR T cell itself, the CAR T cell expresses shRNAs against the HIV 5’LTR and CCR5(27). We have previously shown that our CAR T cells can specifically kill cells that express viral proteins *in vitro*, and can delay viral rebound upon ART cessation in humanized mice that endogenously express these CAR T cells(27,28). The novel combination of SUW133-mediated latency reversal and CAR T cells in depleting latently infected cells *in vitro* has not been investigated to date.

Hence in this study we evaluated whether latently infected cells that are treated with a highly potent, designed LRA (SUW133) can express viral antigens in a way that facilitates their recognition and clearance by CAR T cells. SUW133 is a synthetic analog of bryostatin-1, that has shown superior *in vivo* tolerability relative to the parent compound, as well as improved latency reversal of HIV-infected cells(6,24,25). We have found that *in vitro* generated CAR T cells recognize latently infected U1 cells after SUW133 treatment, as shown by an upregulation of CD107a **(Fig. 5A and Fig. 5B)**. In addition to recognition of SUW133-treated U1 cells, CAR T cells specifically kill SUW133-treated U1 cells **(Fig. 6A-C)**. These data indicate that CAR T cells can recognize and preferentially kill latently infected cells treated with SUW133 *in vitro*.

Because SUW133 can be safely administered *in vivo*, and has been shown to delay viral rebound in humanized mouse models in the absence of CAR T cells(24,25), these *in vitro* data suggest the potential for the combination of this LRA and CAR T cells to further deplete infected cells by reactivating and killing latent HIV reservoir cells *in vivo*. Subsequent studies will examine this possibility.

Naturally-occurring cytotoxic T cells and CAR T cells can each be subject to immune exhaustion during prolonged stimulation(4,28,31). We have shown that CAR T cells that have been exhausted *in vitro* co-express higher levels of Tim-3 and PD-1, which represents terminal exhaustion. However, when CAR T cells are *in vitro* exhausted in the presence of rapamycin, the frequency of Tim-3^+^PD-1^+^ cells decreases significantly **(Fig. 7A-C)**. This same pattern is observed with CD69^+^PD-1^+^ cells, which are also considered to be terminally exhausted **(Fig. S2)**. In rapamycin-treated CAR T cells, CD69 remains upregulated despite a decrease in immune exhaustion markers, indicating continued CAR T cell immunological activation and function.

Together in these studies, we have shown that rapamycin is compatible with *in vitro* latency reversal in J-Lat 10.6 and A2 clones using three distinct classes of LRAs and does not interfere with early T cell activation in human PBMCs. We have also shown that SUW133-treated latently infected cells are recognized and killed by CAR T cells, and that fewer terminally exhausted CAR T cells are present when treated with rapamycin during chronic *in vitro* stimulation. Overall, these data demonstrate the utility of LRAs and rapamycin in the context of a CAR T cell-mediated “Kick-and-Kill” HIV cure approach and provide key insight into achieving a functional HIV cure.

## METHODS

### Isolation of Human Peripheral Blood Mononuclear Cells (PBMCs)

Peripheral blood from HIV seronegative donors was collected by the UCI Institute for Clinical and Translational Science (ICTS) using procedures approved by the UCI institutional review board (IRB), and were provided for this study in a de-identified manner. PBMCs were isolated via Ficoll-Paque (Cat#: 45-001-751, Cytvia) separation and frozen in Bambanker Freezing Media (Cat#: CS-02-002/BB05, NIPPON Genetics) for later use. For all assays, PBMCs were thawed and rested in RF10 (Roswell Park Memorial Institute Medium (RPMI), 10% Fetal Bovine Serum (FBS) 1% Penicillin/Streptomycin) media for 1-2 days before experimentation.

### PBMC Proliferation Assay

PBMCs were co-stimulated for 4 days with Dynabeads Human T-Activator CD3/CD28 (Cat#: 11161D, ThermoFisher Scientific) in the presence of 30 IU/mL IL-2 and 5 nM or 5000 nM rapamycin (Cat#: S1039, Selleck Chemicals) in tissue culture treated round bottom 96 well plates. PBMC cell counts were quantified on a DeNovix CellDrop FL digitized cell counter via acridine orange (Cat# C755G11, Thomas Scientific LLC)/propidium iodide stain (Cat# C755G09, Thomas Scientific LLC).

### PBMC Assessment of CD69 and CD25 Expression Upon Co-Stimulation or Bryostatin-1/Rapamycin Treatment

PBMCs were co-stimulated in the presence of 30 IU/mL IL-2 and treated with 5 nM or 5000 nM rapamycin as described above. PBMCs were also treated with 10 nM or 50 nM bryostatin-1 in the presence/absence of 5 nM and 5000 nM rapamycin. PBMCs were collected and stained for 20 minutes at 4°C with anti-CD3 PerCP/Cy5.5 (Cat#: 300328, BioLegend), anti-CD4 PE (Cat#: 317410, BioLegend), anti-CD8 FITC (Cat#: 344704, BioLegend), anti-CD25 APC (Cat#: 302609, BioLegend), anti-CD69 PE-Cy7 (Cat#: 310912, BioLegend) in 50% Human AB serum (Cat#: HS-20, Fisher Scientific). Cells were stained for 20 minutes at 4°C, and washed with 2% FBS (Cat#: FB-02, Fisher Scientific). Cells were fixed at 4°C overnight in 2% paraformaldehyde (PFA) before flow cytometric analysis (Cat#: 04018-1, Poly Sciences). Collection of flow cytometric data was completed using a BD Fortessa X20, and analysis was completed using FlowJo version 10.9.0 analytical software.

### Quantification of In Vitro HIV Latency Reversal in J-Lat 10.6 and A2 Clones

The following reagents were obtained through the NIH HIV Reagent Program, Division of AIDS, NIAID, NIH: J-Lat Full Length Cells (10.6), ARP-9849, and J-Lat Tat-GFP Cells (A2), ARP-9854, both of which were contributed by Dr. Eric Verdin. J-Lat clones 10.6 and A2 were cultured in RF10 media in the presence of compounds for two days in tissue culture-treated round-bottom 96 well plates. After two days of compound treatment, cells were washed in 2% FBS and fixed in 2% PFA overnight. GFP expression in J-Lat cell lines was quantified via flow cytometry.

### D1D2CAR 4-1BB Lentiviral Vector Production and titer determination

HEK293FT cells were cultured in T875 flasks and transfected with CAR vectors pccLMND-D1D2-41BB-GFP-WPRE-CCR5sh, pMDGL, Rev, and Cocal via a calcium phosphate based transfection. After 3 days of incubation, supernatant was collected and ultracentrifuged at 30,000 RPM for 1.5 hrs at 4°C. Supernatant was aspirated and the virus pellet was resuspended in cold 1x phosphate-buffered saline (PBS) and frozen at - 80°C. D1D2CAR 4-1BB lentiviral vector titer was determined by infection of CEM cells. CEM cells were infected for 3 hours at 37°C while rotating. Post infection, cells were incubated for 2 days at 37°C, and fixed in 2% PFA. GFP expression in CEM cells was quantified via flow cytometry and used to determine viral titer.

### In Vitro CAR T Cell Generation

Bulk human PBMCs from HIV seronegative donors were thawed and cultured in T75 flasks containing RF10 overnight. CD8^+^ T cells were isolated from the bulk PBMCs using a QuadroMACS Separator (Cat#: 130-090-976, Miltenyi Biotec), LS columns (Cat#: 130-042-401, Miltenyi Biotec) and CD8 human microbeads (Cat#: 130-045-201, Miltenyi Biotec) according to the manufacturer’s instructions. Isolated CD8^+^ T cells were co-stimulated with human T Cell TransAct (Cat#: 130-111-160, Miltenyi Biotec) for 2 days in media containing X-VIVO 15 (Cat#: 02-060Q, Lonza), 10% FBS, 1% Penicillin/Streptomycin/Glutamine (Cat#: 10378016, ThermoFisher Scientific) and 50 IU/mL IL-2 (Cat#: LS20002500UG, Gibco) (Growth Media). Co-stimulated CD8^+^ T cells were transduced with D1D2CAR 4-1BB lentiviral vector (MOI 11.5) on RetroNectin (Cat#: T100A, Takara)-coated plates (Not tissue culture treated) via a previously described spin infection protocol(68). Transduction efficiency was determined by GFP^+^ (%) CD8^+^ T cells.

### CAR T Cell In Vitro Exhaustion Assay

24-well tissue culture treated plates were coated with 50,000 ng/mL goat anti-mouse (GAM) IgG secondary antibody (Cat#: 31160, Invitrogen) and 5000 ng/mL anti-human CD3 antibody (Cat#: 70-0037-U100, Tonbo Biosciences). CAR T cells were plated on coated wells and 2000 ng/mL anti-human CD28 antibody added to the growth media (Cat#: 40-0289-U500, Tonbo Biosciences). Cells were incubated for 2 days at 37°C before being transferred to a new well coated with GAM and anti-human CD3 antibody without anti-human CD28 antibody present as described in(58). This was repeated 2 more times, each with a 2-day incubation period. Cells were stained for 20 minutes at 4°C with anti-CD3 PerCP/Cyanine5.5 (Cat#: 300328, BioLegend), anti-CD366 (Tim-3) APC/Fire 750 (Cat#: 345044, BioLegend), anti-CD4 PE (Cat#: 300550, BioLegend), anti-CD8 Spark UV 387 (Cat#: 344776, BioLegend), anti-CD69 (Cat#: 310912, BioLegend), anti-PD-1 (Cat#: 367424, BioLegend), anti-CD25 (Cat#: 302610, BioLegend), and Zombie Violet (Cat#: 423113, BioLegend). After staining cells were fixed in 2% PFA before flow cytometric analysis.

### CAR T Cell CD107a Assay

The following reagent was obtained through the NIH HIV Reagent Program, Division of AIDS, NIAID, NIH: Human Immunodeficiency Virus Type 1 (HIV-1) Infected U937 Cells (U1), ARP-165, contributed by Dr. Thomas Folks. U1 cells were treated with 50 nM SUW133 or mock and incubated for 2 days at 37°C. After this incubation, U1 cells were stained with 1:1000 CellTrace Far Red (Cat#: C34572, Life Technologies) and washed with 2% FBS. CAR T cells were mixed with CellTrace Far Red stained, SUW133/mock treated U1 cells at a 3:1 E:T ratio in a tissue culture treated 96 well plate. Immediately after mixing CAR T cells and U1 cells, 1 uL of anti-CD107a PE (Cat#: 328608, BioLegend) was added directly into each well. After 1 hour, 5 uL of a GolgiStop (Cat#: BDB554724, Fisher Scientific) mix (created by mixing 4 uL of GolgiStop to 150 uL of growth media) was added directly into each well. The plate was incubated for 3 more hours at 37°C. After the 3 hours, the contents of each well were transferred to 1.5 mL Eppendorf tubes and washed twice in 2% FBS. The cells were then stained with 1:1000 Zombie UV (Cat#: 423108, BioLegend) for 20 minutes at room temperature, washed with 2% FBS, and fixed in 2% PFA at 4 C overnight. CD107a expression on CAR T cells was quantified via flow cytometric analysis.

### CAR T Cell Killing Assay

E:T mix of CAR T cells and U1 cells was performed as described in the “CAR T Cell CD107a Assay”. The plate was incubated at 37°C for ∼16 hours. After this incubation, supernatants were collected and stored at -80°C, and the cells were stained with 1:1000 Zombie UV for 20 minutes at room temperature. Cells were permeabilized and fixed using a BD Cytofix/Cytoperm Fixation/Permeabilization Solution Kit (Cat#: BDB554714, Fisher Scientific) and intracellularly stained with KC57-RD1 (Cat#: 6604667, Beckman Coulter). Live Gag+ U1 cells were quantified via flow cytometry. Specific killing was measured by (percent live Gag^+^ U1 cells without CAR T cells – percent live Gag^+^ U1 cells with CAR T cells)/(percent live Gag^+^ U1 cells without CAR T cells). Specific killing of mock treated U1 cells was calculated using %live U1 cells instead of % live Gag^+^ U1 cells, as mock treated U1 cells do not express gag.

### Synthesis and Structure of SUW133

SUW133 was prepared using the previously reported Bryostatin-1 analog synthesis developed by DeChristopher et al., which relies on a convergent fragment-coupling approach, and a Prins-driven macrocyclization(69).

## Author Contributions

CRediT: **Nishad S Maggirwar**: Conceptualization, Data curation, Formal Analysis, Funding acquisition, Investigation, Methodology, Project administration, Validation, Visualization, Writing – original draft, Writing – review & editing; **José A Morán**: Data curation, Funding acquisition, Investigation, Validation, Visualization, Writing – review & editing; **Wenli Mu**: Conceptualization, Methodology, Validation, Visualization, Writing – review & editing; **Thomas D Zaikos**: Conceptualization, Methodology, Visualization, Writing – review & editing; **Tessa Chou**: Visualization, Writing – original draft, Writing – review & editing; **Shireen R Turner**: Formal Analysis, Investigation, Writing – review & editing; **Brian H Yu**: Formal Analysis, Investigation, Writing – review & editing; **Alok Ranjan:** Conceptualization, Methodology; **Rami Hourani:** Conceptualization, Methodology; **Paul A Wender**: Funding acquisition, Supervision, Writing – review & editing; **Anjie Zhen**: Conceptualization, Funding acquisition, Methodology, Supervision, Writing – review & editing; **Matthew D Marsden**: Conceptualization, Funding acquisition, Resources, Project administration, Supervision, Writing – review & editing

## Funding

The authors’ laboratory is funded by the National Institute of Health grant number AI172727 (A.Z. and M.D.M.). The design, synthesis and preliminary binding and cellular activity of SUW133 was funded by National Institute of Health grants to P.A.W. (R01CA031845 and AI172410). J.A.M. is a predoctoral trainee supported by U.S. Public Health Service training grant T32 AI007319 from the NIH. N.S.M is a predoctoral trainee supported by an Individual National Research Service Award (F31). Research reported in this publication was supported by the National Institute of Allergy and Infectious Diseases of the National Institutes of Health under Award Number F31AI192129. The content is solely the responsibility of the authors and does not necessarily represent the official views of the National Institutes of Health.

## Acknowledgments

Figures were created using Biorender.com.

## Conflicts of Interest

Stanford University has filed patents on bryostatin and other PKC modulators which have been licensed by Neurotrope BioScience for the treatment of neurological disorders and by Bryologyx Inc. for use in HIV/AIDS eradication and cancer immunotherapy. P.A.W is an advisor to both companies and a cofounder of the latter.

