## Supplemental Figures for "Leveraging HIV-Specific CAR T Cells and Rapamycin Treatment in “Kick-and-Kill” HIV Cure Approaches"

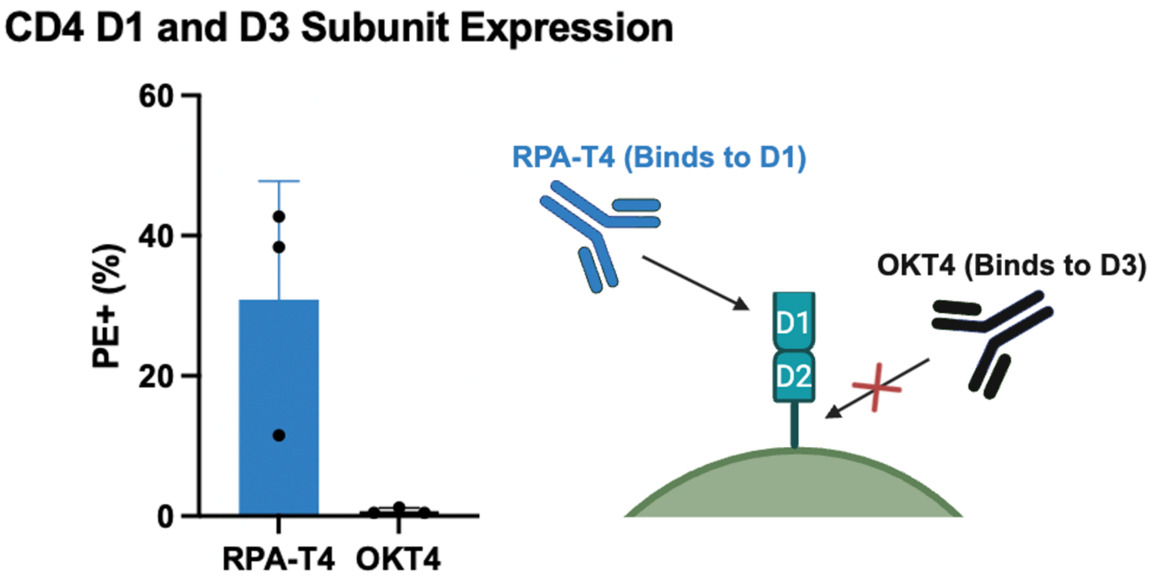


**Fig. S1 Absence of the D3 subunit of CD4 in transduced HEK293FT cells.** HEK293FT cells were cultured for 24 hours before transduction with a D1D2CAR 4-1BB-encoding lentiviral vector. HEK293FT cells were stained with anti-CD4 antibody clones RPA-T4 and OKT4. RPA-T4 is specific for the D1 subunit of CD4, while OKT4 is specific for the D3 subunit. D1 and D3 expression was quantified by $PE^+$ (%) cells acquired via flow cytometry.

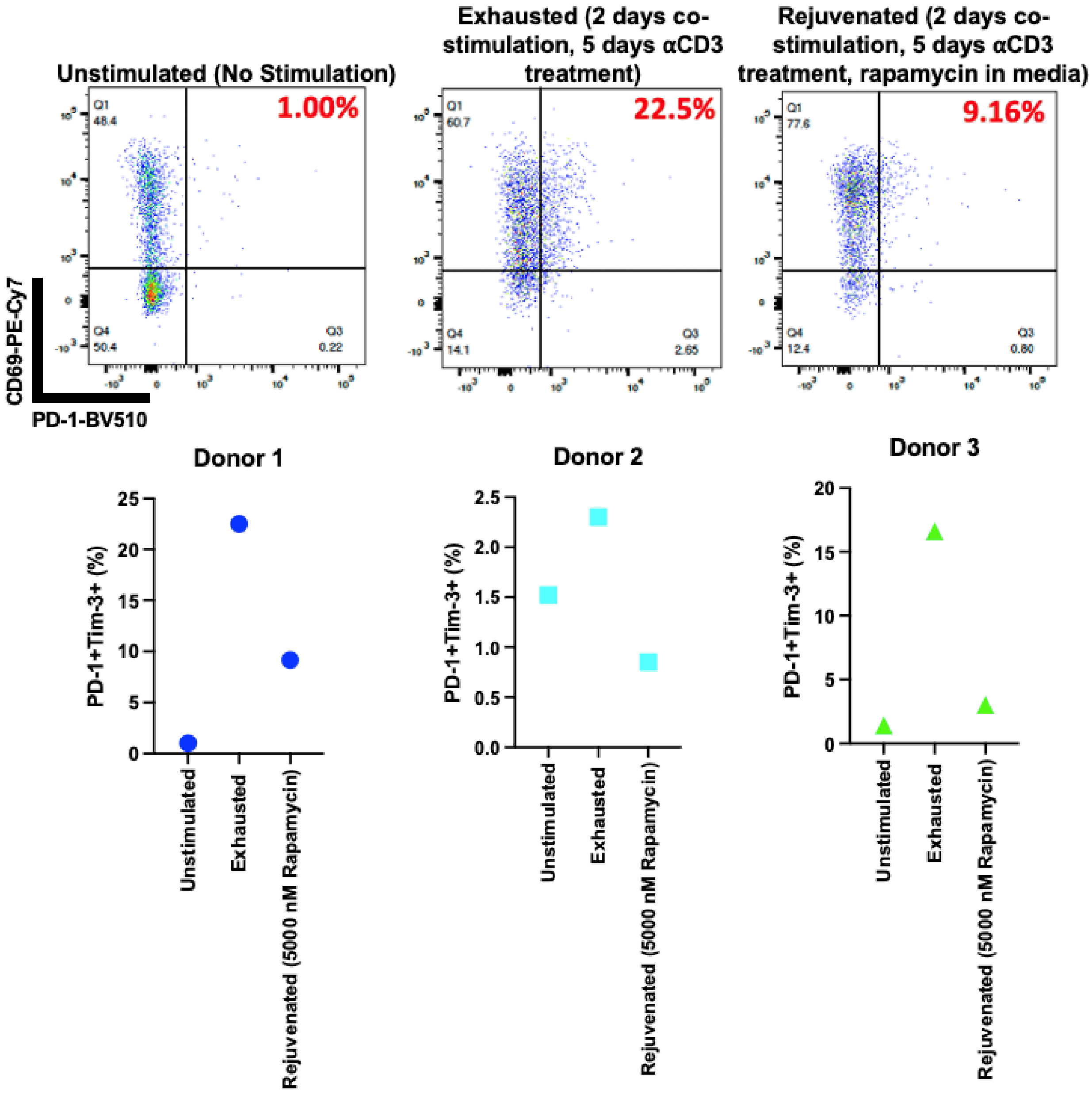


**Fig. S2 CD69+PD-1+ cells are less abundant in chronically stimulated, rapamycin treated, D1D2CAR T cells.** D1D2CAR T cells were generated as described in Fig. 8. $CD69^{+}PD\text{-}1^{+}$ cells were quantified via flow cytometry. N = 3 unique HIV seronegative human PBMC donors (indicated by a unique colored symbol).

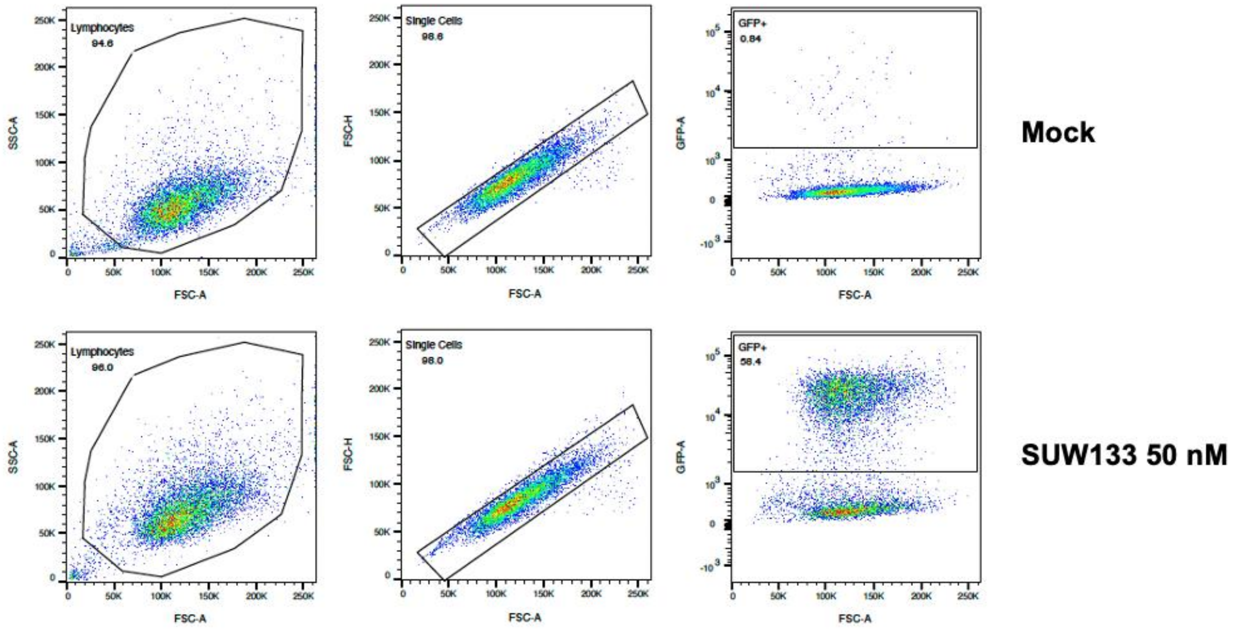


**Fig. S3 J-Lat clone gating strategy.** The J-Lat clone population was identified via FSC-A/SSC-A. Doublet exclusion was performed before identifying the $GFP^+$ cell population. $GFP^+$ gate was determined using the mock control.

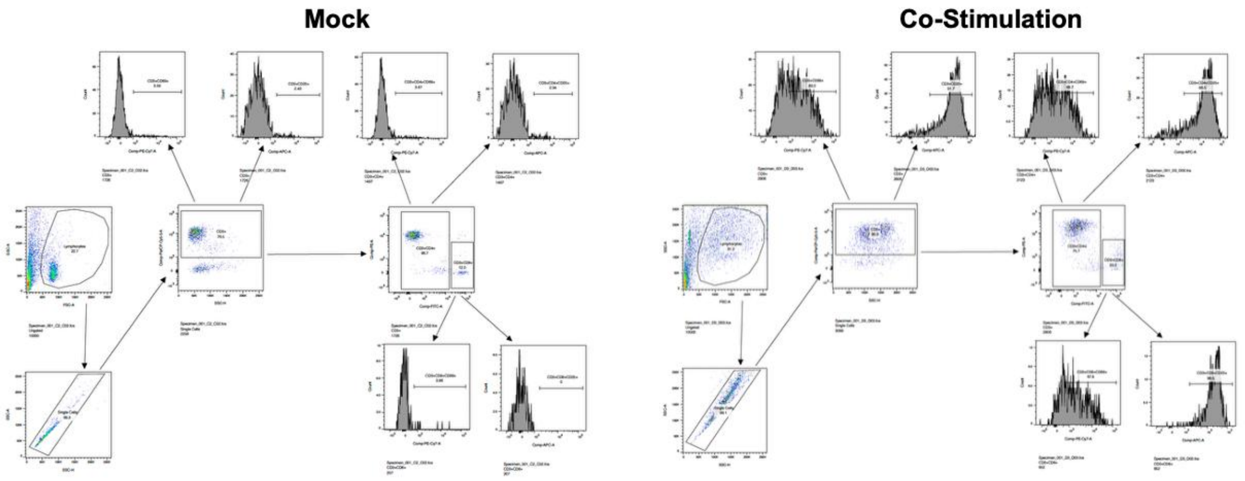


**Fig. S4 PBMC Stimulation Gating Strategy.** Total lymphocytes were identified via FSC-A/SSC-A. Doublet exclusion was performed before identifying the $CD3^+$ population. CD69 and CD25 expression was then assessed from the $CD3^+$ population. $CD4^+$ and $CD8^+$ cell populations were identified from the $CD3^+$ cell population. CD69 and CD25 expression was assessed in the $CD4^+$ and $CD8^+$ cell populations. Gating was determined based on isotype controls.
